# Measuring the host-seeking ability of *Aedes aegypti* destined for field release

**DOI:** 10.1101/695528

**Authors:** Meng-Jia Lau, Nancy M. Endersby-Harshman, Jason K. Axford, Scott A. Ritchie, Ary A. Hoffmann, Perran A. Ross

## Abstract

Host-seeking is an essential process in mosquito reproduction. Field releases of modified mosquitoes for population transformation rely on successful host-seeking by female mosquitoes, but host-seeking ability is rarely tested in a realistic context. We tested the host-seeking ability of female *Aedes aegypti* mosquitoes using a semi-field system. Females with different *Wolbachia* infection types (*w*Mel-, *w*AlbB-infected and uninfected) or from different origins (laboratory and field) were released at one end of a semi-field cage and recaptured as they landed on human experimenters fifteen meters away. Mosquitoes from each population were then identified with molecular tools or through marking with a consistent weight of fluorescent powder. *Wolbachia*-infected and uninfected populations had similar average durations to landing and overall recapture proportions, as did laboratory and field-sourced *A. aegypti*. These results suggest that the host-seeking ability of mosquitoes is not negatively affected by *Wolbachia* infection or long-term laboratory maintenance. This method provides an approach to study the host-seeking ability of mosquitoes across a long distance which will be useful when evaluating strains of mosquitoes that are planned for releases into the field to suppress arbovirus transmission. An adjustment of this method may also be useful in sterile insect release programs because male host-seeking and swarming around female feeding sites can also be investigated.

## Introduction

The management of arboviral diseases has become increasingly important to global health in recent decades.^1^ The occurrence of arboviral diseases such as dengue, Zika, Japanese encephalitis and West Nile fever is increasing, especially in tropical and subtropical areas. ^2, 3, 4^ These viruses require blood-feeding mosquitoes to complete their life cycle,^5^ with mosquitoes from the genera of *Culex* and *Aedes* being particularly important.^6^ An effective way to control arbovirus transmission is to suppress the vector mosquito populations. Pesticides are widely used for this purpose but this can lead to the evolution of physiological resistance, alongside other undesirable effects associated with pesticide use.^7, 8^ The sterile insect technique (SIT),^9^ incompatible insect technique (IIT),^10^ and the release of insects carrying a dominant lethal gene (RIDL)^11^ are promising non-insecticidal alternatives, where wild-type females that mate with the released “modified” males have few viable offspring, decreasing the population size.

An alternative approach aims to decrease the ability of mosquitoes to transmit viruses by introducing endosymbiotic *Wolbachia* bacteria.^12, 13^ *Wolbachia* are transmitted maternally and can invade natural populations through cytoplasmic incompatibility and any beneficial effects on host reproduction.^14, 15^ When introduced into mosquitoes from other insects, some *Wolbachia* strains reduce their capacity to transmit viruses.^12, 16^ *Aedes aegypti* infected with the *w*Mel *Wolbachia* strain have been introduced into field populations, with the first releases taking place in Cairns, Australia in 2011.^17^ In locations in Australia where *Wolbachia* have established there have been no confirmed locally-transmitted cases of dengue occurring within the release areas.^18, 19^

Population replacement and suppression strategies ideally should be preceded by investigations to assess their potential for success, address safety concerns,^20^ and perform community engagement.^18, 21^ When using *Wolbachia* to block arbovirus transmission, fitness costs imposed on their hosts such as adult life-shortening,^22^ reduced quiescent egg viability,^23^ and reduced starvation resistance of larvae^24^ must be considered. Such effects mean that *Wolbachia* must exceed a threshold frequency in order to spread in natural populations.^17, 25, 26^ SIT, IIT and RIDL programs are simpler in that the only concern is male fitness, but still require the released males to have a high competitiveness to ensure successful mating with wild females.^27^ Populations reared in the laboratory can adapt to the artificial conditions which may reduce field performance.^28, 29^ For instance, laboratory maintenance can lead to the loss of pesticide resistance,^30^ greatly reducing fitness in release areas with heavy pesticide use.^21^

Fitness assays are usually carried out in the laboratory to detect fitness costs, but during releases mosquitoes must locate hosts or mates under variable environmental conditions. Performance under laboratory conditions often does not translate to performance in the field.^31, 32, 33^ Males from the transgenic OX3604C strain of *A. aegypti* successfully suppressed laboratory populations^34^ but were much less effective under semi-field conditions due to a strong mating disadvantage.^35^ For *Wolbachia* releases, density-dependent effects,^25^ loss of cytoplasmic incompatibility,^36^ and incomplete maternal transmission^37^ may account for the slower-than-expected spatial spread of infections in natural populations^37, 38^ or even failed establishment^39^ despite success under more controlled conditions.^12^

Successful host-seeking is key to population replacement programs since female mosquitoes require blood for reproduction. Females locate a potential blood source by tracking exhaled CO_2_ over tens of meters, then approach and land on the host by detecting thermal plumes, host odors, moisture and visual contrast.^40, 41, 42^ *Wolbachia* infections do not affect the attraction of *A. aegypti* to human odors in the laboratory,^43^ but successful host-seeking in the field will depend on the detection of olfactory cues from a long distance, visual and temperature signals from a shorter distance and flight ability.

In this paper, we tested the host-seeking ability of female *A. aegypti* using a semi-field cage in North Queensland, Australia^44^ to simulate an outdoor setting. Females were released at one end of the semi-field cage and then recaptured by two experimenters seated at the other end. This method allows for a direct comparison of host-seeking ability between different mosquito strains in a common environment. To test the method we compared mosquitoes with the *w*Mel and *w*AlbB *Wolbachia* strains, which are now being released into the field in disease control programs,^17^ (Nazni et al., unpublished data) against uninfected counterparts. To evaluate whether laboratory adaptation could affect host-seeking as demonstrated in laboratory experiments previously,^45^ we also compared a laboratory population to a population collected recently from the field.

## Material and Methods

### Mosquito strains and maintenance

*A. aegypti* mosquitoes in this study were reared at 26-28°C in a controlled temperature room at James Cook University, Cairns, using methods described previously.^46^ We performed two sets of experiments to compare the effects of *Wolbachia* infection and laboratory maintenance on host-seeking ability respectively. To test for the effects of *Wolbachia* infection, we used uninfected, *w*Mel-infected and *w*AlbB-infected *A. aegypti* with a similar genetic background. Populations infected with *w*Mel and *w*AlbB were derived from lines transinfected previously.^12, 47^ The *w*Mel population was collected from Cairns, Australia in May 2013 from regions that had been invaded two years earlier^17, 48^ while the *w*AlbB population was crossed to an Australian background and maintained in the laboratory.^49^ The uninfected population was established from *A. aegypti* (*Wolbachia*-uninfected) eggs collected in Cairns, Queensland, Australia in November 2015.^50^ Females from all *Wolbachia*-infected lines were backcrossed for three generations to the uninfected males to ensure a similar genetic background before the experiments.^23^ To test for the effects of laboratory maintenance we compared the host-seeking ability of laboratory and field populations. The laboratory population was identical to the uninfected population described above and had been maintained in the laboratory for 27 generations. The field population of *A. aegypti* (*Wolbachia*-uninfected) was collected in September 2018 from the same location as the laboratory population and was a mix of the first and second laboratory generations at the time of experiments.

For each release, the compared colonies were hatched synchronously, provided with TetraMin^®^ fish food tablets (Tetra, Melle, Germany) *ad libitum* and the larval density was controlled to 150 in 1 L water to ensure matched eclosion. After pupation, approximately 80 pupae were selected with a mix of 80% females and 20% males and left to emerge as adults in one cage (BugDorm-4M1515 Insect Rearing Cage). Each cage was provided with a cup of 10% sucrose and water and left for at least 4 d to ensure that females had matured and mated, but not blood fed. One day before the release, sugar cups were removed with only water cups remaining to starve the females, since sugar feeding may affect host-seeking behavior.^51, 52^ The released females were 5 d old in both the *Wolbachia* infection comparison and the laboratory maintenance comparison.

### Release-recapture method

We used a semi-field system (17.5 × 8.4 m) at James Cook University, Cairns, Australia containing soil, vegetation, a “Queenslander” house structure (Qld) and a ventilation system to match outside ambient temperatures to simulate natural conditions (Figure 1).^44^ Mosquitoes were released near the door side from a box with a mesh lid while two experimenters were seated within the Qld structure to attract mosquitoes from the other end (Figure 1). Two temperature loggers (Thermochron; 1-Wire, iButton.com, Dallas Semiconductors, Sunnyvale, CA, USA) were placed near the release point and two were placed under the Qld structure to monitor temperatures during experiments (Supplementary Table 1).

**Figure 1.**
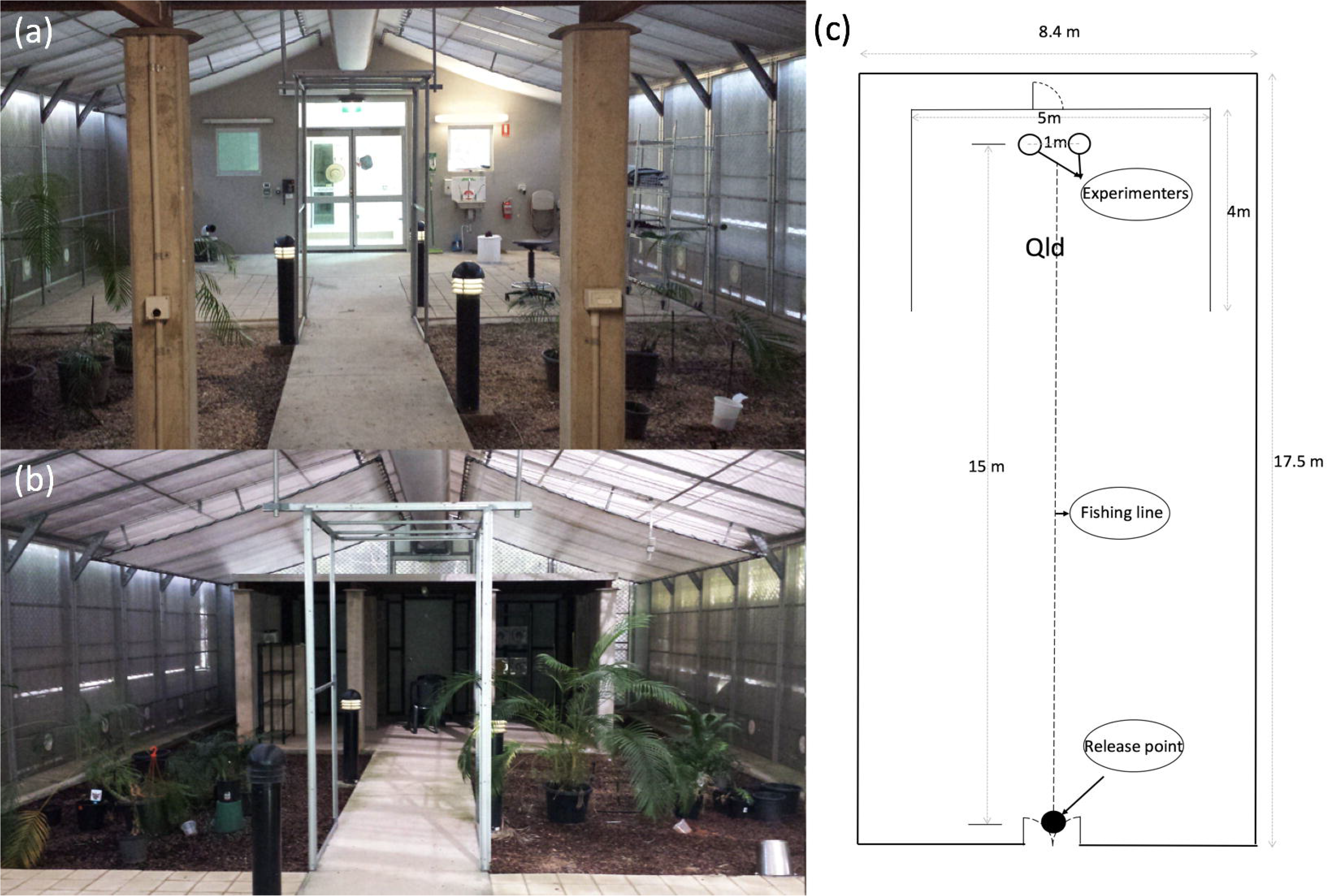
Interior of the semi-field cage. (a) View of the door from inside the Qld. (b) View of the Qld from the door. (c) Schematic diagram of the cage showing the release point and the location of two experimenters.

Females from all populations in the comparison were aspirated into a single release box (Supplementary Figure 1) and placed in the semi-field cage to acclimate for at least 30 minutes before experiments commenced. For the *Wolbachia* infection comparison, 50 uninfected, 50 *w*Mel-infected and 50 *w*AlbB-infected females were released into the box. For the laboratory maintenance comparison, 50 laboratory and 50 field source females were released. Females that were damaged during handling were replaced.

Two experimenters wore bug net mesh hats, long-sleeved shirts and shorts, exposing only their lower legs to restrict the area where mosquitoes could land. The same two experimenters undertook all experiments. Experimenters sat on the floor within the Qld structure, 1 m apart (Figure 1) with an electronic timer, mechanical aspirators (Model 2809C, BioQuip Products, Inc, Rancho Dominguez, CA, USA) and 15 collection vials nearby. The experiment commenced by pulling the fishing line to remove the mesh lid from the box to release the mosquitoes (Supplementary Figure 1), after which the timer was immediately started. Females landing on exposed skin were collected with mechanical aspirators as they landed. Collection vials were replaced with empty vials at 3-minute intervals until 42 minutes had elapsed. After 42 minutes, both experimenters moved to the opposite end of the cage to capture mosquitoes that did not land during the experiment. Collections occurred until no more mosquitoes were detected after a thorough search of the semi-field cage. Between experiments, two Biogents Sentinel (BGS) traps (Biogents AG, Regensburg, Germany) were placed inside the semi-field cage to assist in the capture of any remaining mosquitoes. At least one hour before each experiment commenced, the experimenters searched the semi-field cage and used an electric mosquito swatter to kill any mosquitoes found.

### Wolbachia infection comparison

The host-seeking experiment was repeated seven times with 50 uninfected, 50 *w*Mel-infected and 50 *w*AlbB-infected females. Females collected from each replicate and time interval were stored in absolute ethanol at 4°C for wing length measurements, DNA extraction and *Wolbachia* screening. One replicate was discarded from other analyses due to the loss of samples during wing dissection.

Field-collected *A. aegypti* are smaller and more variable in size than laboratory-reared *A. aegypti*.^53^ Since host-seeking females collected from the field in a previous experiment tended to be larger than non-host-seeking females,^54^ we tested whether host-seeking speed and successful host seeking within 42 minutes was affected by size. We measured the wing length of females from two experimental replicates to obtain an indication of their body size.^55^ Intact wings were dissected from individual females and fixed under a 10 mm circular coverslip (Menzel-Gläser, Braunschweig, Germany) using Hoyer’s solution^56^ for further observation and measurement with an NIS Elements BR imaging microscope (Nikon Instruments, Japan).^24^

DNA extraction and *Wolbachia* screening were conducted according to the methods of Lee, et al. ^57^ DNA from whole mosquitoes was extracted using 200 µL of 5% Chelex 100 Resin (Bio-Rad Laboratories, Hercules, CA) and 3 μL of Proteinase K (20 mg/ mL, Bioline Australia Pty Ltd, Alexandria NSW, Australia). Extractions were diluted by 1/10, pipetted into four positions of a 384-well plate and amplified with mosquito-specific (*mRpS6*) primers, *A. aegypti*-specific (*aRpS6*) primers, *Wolbachia w*Mel-specific (w1) primers and *Wolbachia w*AlbB-specific (*wAlbB*) primers^48, 49, 58, 59^ using a LightCycler 480 system (Roche Applied Science, Indianapolis, IN, USA). Robust and similar amplification of *mRpS6* and *aRpS6* (within one cycle) was expected for each individual. Uninfected *A. aegypti* were expected to show no amplification and therefore, no crossing point (Cp) value, with both *w1* and *wAlbB* primers. *A. aegypti* were classified as *w*Mel-infected when they exhibited no amplification with *wAlbB* primers and low Cp values (< 28) and a Tm within the expected range for *w1* primers based on *w*Mel-infected laboratory controls. *w*AlbB-infected *A. aegypti* tested positive for *wAlbB*, *mRpS6* and *aRpS6* primers but also showed late amplification (Cp > 28) with *w1* primers. Individuals were therefore classified as *w*AlbB-infected when they exhibited a low Cp value (< 28) with *wAlbB* primers, a Tm within the expected range for *wAlbB* primers and an amplification curve shape consistent with *w*AlbB-infected laboratory control values (Supplementary Figure 2). At least two consistent technical replicates were obtained for each individual.

### Laboratory maintenance comparison

In this experiment, laboratory and field populations were marked with different colors of fluorescent powder (DayGlo, Barnes Products Pty Ltd, Moorebank, NSW, Australia) before release since the two populations could not be distinguished by molecular assays. Orange, blue and yellow colors were used and were cycled between replicates. To reduce potential negative effects of marking, we used a minimal, but visually identifiable amount (Supplementary Figure 3) by weighing powder on a microbalance (Sartorius BP 210 D). One hour before the release, 50 females from each population were aspirated into two separate 70-mL specimen cups containing approximately 0.4 mg of fluorescent powder in different colors. The cups were shaken gently to coat the mosquitoes evenly in powder before placing them in the release box (Supplementary Figure 1). Recaptured females were killed by freezing at −20 °C for 30 minutes and identified under a microscope using a UV flashlight. This experiment was repeated six times.

### Data analyses

Data visualization and ANOVA analyses were conducted using R studio with the packages Rmisc,^60^ plyr,^61^ and ggplot2.^62^ Mosquitoes were captured at three-minute intervals and assigned a value based on the median time of each catching interval for average landing time calculations. Mosquitoes caught after 42 minutes were considered as not landing. A two-way ANOVA analyzed differences in average landing time of the landed mosquitoes and the number of females that landed by treating population as a fixed factor and experimental replicate as a random factor. One-way ANOVA was used to compare the wing length of mosquitoes caught at different intervals by treating landing time as a factor. Cumulative landing proportions over time were analyzed with log-rank tests in IBM SPSS Statistics version 25 by combining replicate experiments together.

## Results

### Wolbachia infection comparison

We compared the host-seeking ability of uninfected, *w*Mel-infected and *w*AlbB-infected females when released simultaneously in a semi-field cage. On average, more than 30% of the mosquitoes were captured during the first three minutes of the experiment, with approximately 70% landing over the course of 42 minutes (Figure 2A). We compared the cumulative landing proportions of each population when combined across replicates and found no significant differences between *Wolbachia*-infected and uninfected females (log-rank: *w*Mel : uninfected: χ^2^ = 1.428, df = 1, P = 0.232; *w*AlbB : uninfected: χ^2^ = 2.9, df = 1, P = 0.089, Figure 3A).

**Figure 2.**
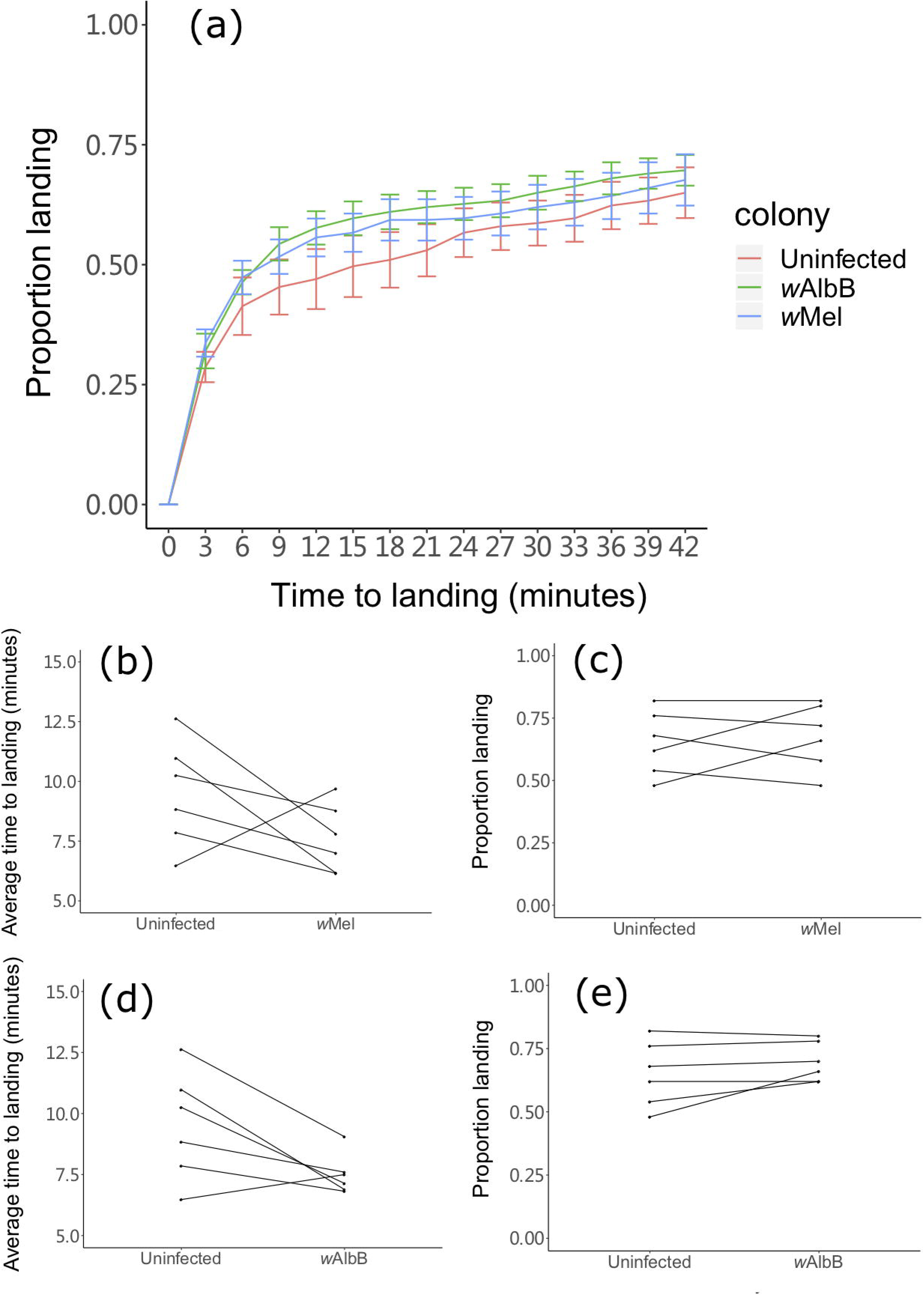
Host-seeking ability of 5 d old *w*Mel-infected, *w*AlbB-infected and uninfected *A. aegypti* females in a semi-field cage. (a) Cumulative landing proportions of females on human experimenters across all replicates. Lines represent means and error bars represent standard errors. (b-c) Comparisons of average time to landing (b) and proportion landing (c) between uninfected and *w*Mel-infected females, plotted separately for each replicate. (d-e) Comparisons of average time to landing (d) and proportion landing (e) between uninfected and *w*AlbB-infected females, plotted separately for each replicate.

**Figure 3.**
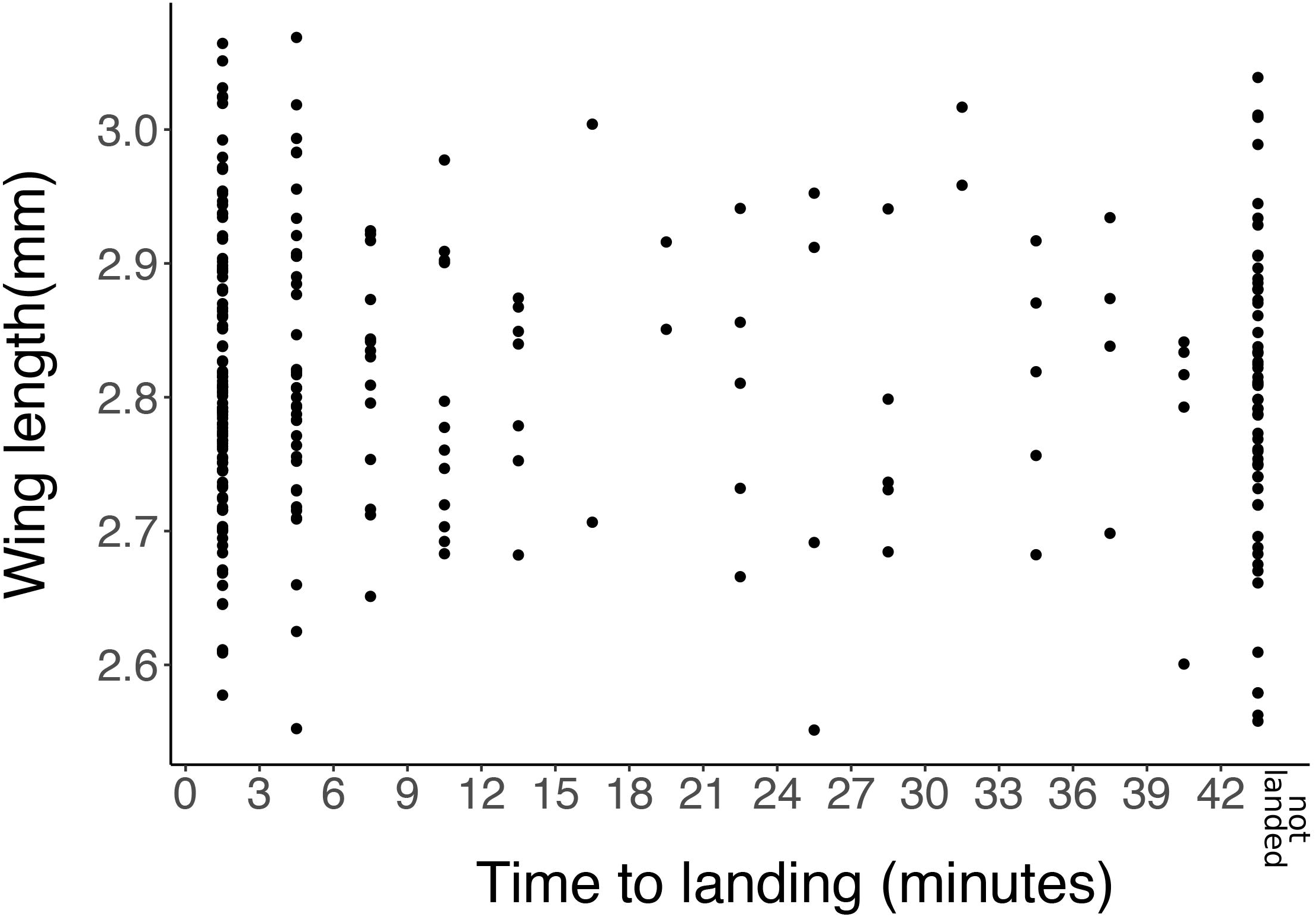
Wing lengths of female *A. aegypti* collected during two replicates of the *Wolbachia* infection host-seeking experiment. Points represent wing lengths of individual females collected across each 3-minute interval of the experiment. Wing lengths of females captured after 42 minutes had elapsed were also included.

The average time to landing of each population was used as an estimate of host-seeking speed (Figure 2b, 2d). Average time to landing did not differ significantly between uninfected (mean ± SE: 9.5 ± 0.9 minutes), *w*Mel-infected (7.6 ± 0.6 minutes) and *w*AlbB-infected (7.5 ± 0.3 minutes) females (two-way ANOVA: *w*Mel : uninfected: F_1,5_ = 2.503, P = 0.174; *w*AlbB : uninfected: F_1,5_ = 6.434, P = 0.052). There was also no significant effect of replicate on average time to landing in either comparison (*w*Mel : uninfected: F_5,5_ = 0.617, P = 0.696; *w*AlbB : uninfected: F_5,5_ = 2.009, P = 0.231). We compared the total proportion of females landing as an indicator of overall host-seeking success (Figure 2c, 2e); here there were also no significant differences between populations (*w*Mel : uninfected: F_1,5_ = 0.282, P = 0.618, *w*AlbB : uninfected: F_1,5_ = 2.426,P = 0.180). There was a significant effect of replicate in the *w*AlbB comparison (F_5,5_ = 7.505, P = 0.023) but not in the *w*Mel comparison (F_5,5_ = 3.482, P = 0.099).

Females from two replicates of the *Wolbachia* infection comparison were measured for wing length (Figure 3). There was no significant effect of wing length on host-seeking speed, measured by capture interval (F_14,251_ = 0.708, P = 0.766). Females landing within the first three minutes (2.81 ± 0.02 mm, n = 102) did not differ in size from females collected after 42 minutes had elapsed (2.80 ± 0.03 mm, n = 57), suggesting no difference in size between fast host-seeking females and non-host-seekers (F_1,80_ = 1.311, P = 0.256).

### Laboratory adaptation comparison

In comparisons of laboratory and field *A. aegypti* females, approximately 60% of the released mosquitoes were caught over the duration of the experiments. Cumulative landing proportions did not differ significantly between field and laboratory populations when combined across replicates (log-rank: χ^2^ = 2.275, df = 1, P = 0.131, Figure 4a). The average time to landing did not differ significantly between field (mean ± SE: 12.3 ± 1.1 minutes) and laboratory (10.5 ± 1.3 minutes) females (two-way ANOVA: F_1,5_ = 2.346, P = 0.186, Figure 4b), with no significant effect of replicate (F_5,5_ = 2.876, P = 0.136). Furthermore, the total proportion of females landing did not differ between field and laboratory females (F_1,5_ = 0.745, P = 0.428, Figure 4c), with no significant effect of replicate (F_5,5_ = 4.647, P = 0.059), suggesting that laboratory maintenance does not affect host-seeking ability.

**Figure 4.**
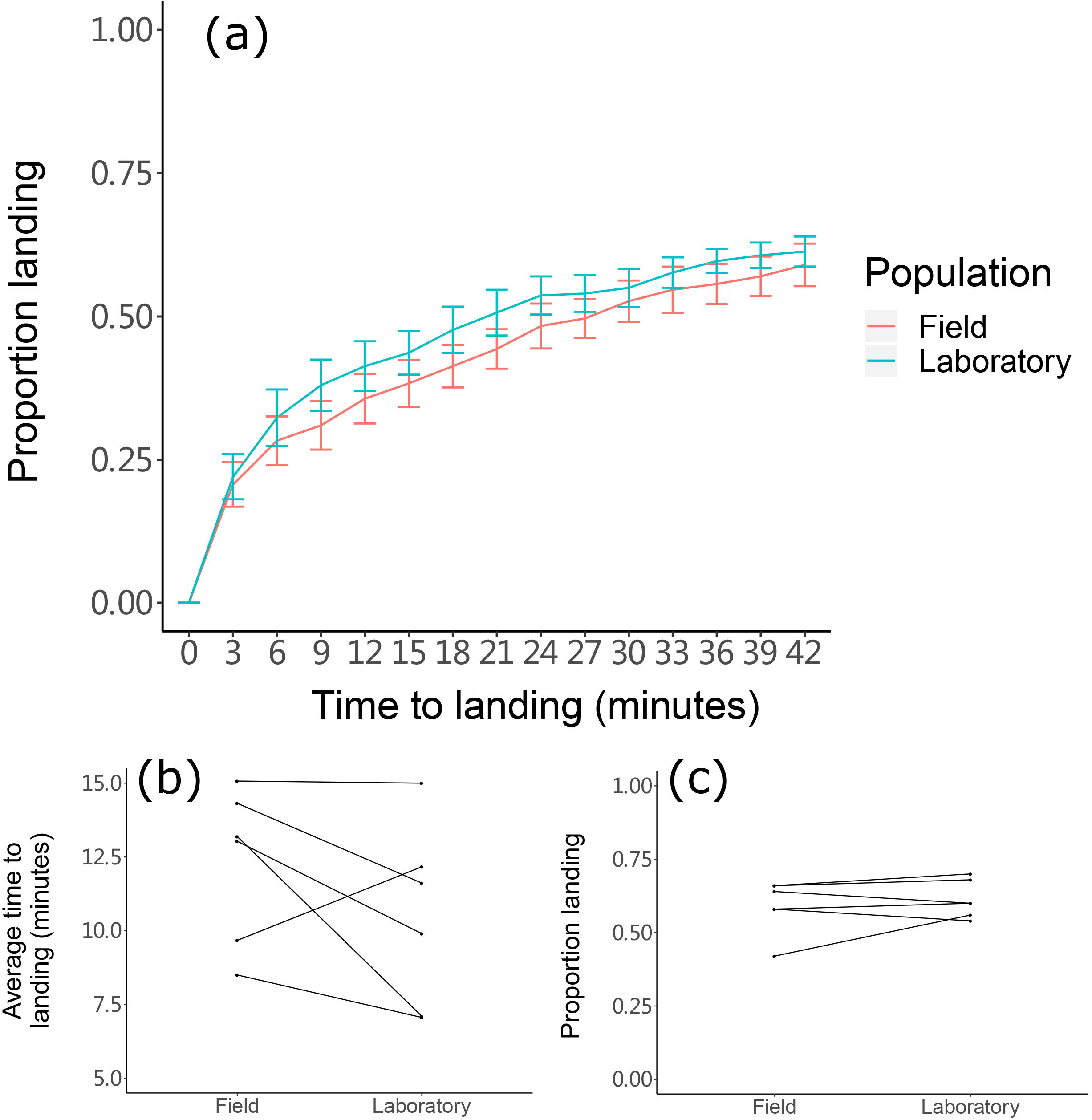
Host-seeking ability of field and laboratory *A. aegypti* females in a semi-field cage. (a) Cumulative landing proportions of females on human experimenters across all replicates. Lines represent means and error bars represent standard errors. (b-c) Comparisons of average time to landing (b) and proportion landing (c) between field and laboratory females, plotted separately for each replicate.

## Discussion

Suppressing the transmission of dengue and other arboviruses by releasing *Wolbachia*-infected mosquitoes is becoming increasingly popular, with releases taking place in at least 12 countries (https://www.worldmosquitoprogram.org/; https://www.nea.gov.sg/corporate-functions/resources/research/wolbachia-aedes-mosquito-suppression-strategy/project-wolbachia-singapore; https://www.imr.gov.my/wolbachia/). For releases to succeed, the strain intended for deployment needs to have comparable fitness to wild-type mosquitoes, which should be tested prior to large-scale field release. The semi-field cage setting is widely used as an intermediate step between laboratory studies and open field releases. ^63, 64, 65^ Semi-field experiments have been used to test the mating success and invasive ability of *Wolbachia* infections^12, 66, 67^ and for evaluating the efficacy of novel mosquito traps and pesticides.^65, 68, 69^ But while host-seeking is critical for the success of *Wolbachia* replacement programs, the strains used in field releases including *w*Mel and *w*AlbB have not been evaluated for their effects on host-seeking ability in a realistic way.

We compared the host-seeking ability of female *A. aegypti* with different *Wolbachia* infection types and from laboratory and field origins in a semi-field cage. Our method was similar to the method developed by McMeniman, et al. ^70^ In their study, the host-seeking ability of wild-type and *Gr3* mutant females lacking a response to CO_2_ was compared by releasing mosquitoes in the middle of the cage and leaving them to disperse naturally for 5 hours before the experiment. In our design, female mosquitoes were released simultaneously at a single release point fifteen meters away from the experimenters, thus standardizing the distance over which host-seeking is tested and allowing mosquitoes to combine their flight ability with the detection of olfactory, visual and thermal queues to locate and land on experimenters. This is the first time that a semi-field approach has been used to evaluate the host-seeking ability of mosquitoes with *Wolbachia* strains intended for field deployment.

We found no significant differences between *A. aegypti* with different *Wolbachia* infection types on host-seeking ability in our experiments. Females with the *w*Mel and *w*AlbB strains should therefore not be at a disadvantage in terms of host-seeking if released into the field. Although a study with a Puerto Rican *A. aegypti* population indicated that laboratory maintenance altered attraction to human odors,^45^ no significant differences were found in overall host-seeking between laboratory and field populations in our semi-field experiments. Therefore, our laboratory maintenance protocol^46^ should not lead to compromised host-seeking ability in the field, though other factors that can coincide with laboratory maintenance such as inbreeding may reduce fitness.^71^ Different rearing procedures, such as the use of membrane feeders, non-human blood or small cages may also affect host-seeking ability if adaptation occurs.

In the absence of molecular tools, visual marking is needed to distinguish between populations in the same experiment. However, overapplication of powder may affect longevity and behavioral responses, with effects depending on the method and the color used for marking.^72, 73, 74^ Although the two sets of experiments were conducted at different times, no significant differences were found between marked and unmarked uninfected laboratory females in terms of average arrival time (One-way ANOVA: F_1,10_ = 0.388, P = 0.547) and proportion landing (F_1,10_ = 0.387, P = 0.548), which suggests that the minimal amount of fluorescent powder used for marking does not affect host-seeking ability.

We also ran experiments with *w*Mel, *w*AlbB-infected and uninfected *A. aegypti* females that were 20 d old and found no significant differences between populations (Supplementary Figure 4). Although mosquitoes of different ages were not compared in the same experiment, we found that 20 d old females had slower average times to landing (Two-way ANOVA: ages: F_1,24_ = 8.567, P = 0.007, colonies: F_2,24_ = 1.407, P = 0.264) but higher landing proportions (ages: F_1,24_ = 5.802, P = 0.024, colonies: F_2,24_ = 0.461, P = 0.636) compared to 5 d old females by treating mosquito age and colony as fixed factors. This suggests that host-seeking ability may be influenced by mosquito age, but direct comparisons between ages in the same experiment are needed to confirm this finding.

Many factors can influence mosquito attraction to humans including environmental temperature and humidity, in addition to the CO_2_, skin emanations, body heat and moisture of the host.^75, 76, 77^ While all mosquitoes in each experiment were reared under the same conditions and were a similar age, we observed substantial differences in average times to landing and landing proportions between replicates (Supplementary Tables 2 and 3). We found no effect of temperature or the time of releases (Supplementary Table 1) according to Spearman’s rank correlation (P > 0.05), suggesting that temperature and time of day did not substantially influence host-seeking. We also ran a power analysis using an online calculator (http://powerandsamplesize.com/Calculators/Compare-2-Means/2-Sample-Equality) with a 80% power test using the average times and standard deviations of *Wolbachia*-infected and uninfected colonies. For 20% differences in our studies (7.5 minutes for *w*Mel or *w*AlbB-infected *A. aegypti* while 9.5 minutes for uninfected *A. aegypti*), at least 15 replicates are needed to detect an effect, while a difference of 30% could be detected with six replicates.

In addition to studying the host-seeking ability of females, it may be possible to extend this method to male mosquitos. For SIT, IIT and RIDL approaches, testing the competitiveness of males before the release is essential.^27^ A previous semi-field cage study showed that *Wolbachia* infection does not reduce the competitiveness of *A. aegypti* males.^67^ However, in nature, adult female densities will not be as high as in semi-field cage tests; males will typically locate and fly around a human host first before detecting female flight tones and initiating courtship behaviour.^78, 79, 80^ In a pilot experiment where we released males into the semi-field cage, we found that *A. aegypti* males exhibited a similar host-seeking response to females (Supplementary Figure 5), but this requires further testing.

In conclusion, we have developed a method to test the host-seeking ability of female *A. aegypti* populations under semi-field conditions. While changes in host-seeking behavior due to *Wolbachia* infections and laboratory adaptation are apparent from some laboratory studies, it is important to test host-seeking in a way that reflects natural conditions. Comparisons of host-seeking ability using this approach will be informative when evaluating mosquito strains for field release. This method can also be used to compare other factors such as age and rearing conditions which can help to better understand the host-seeking behavior of female mosquitoes.

## Acknowledgements

We thank Qiong Yang and Ashley Callahan of the Pest and Environmental Adaptation Research Group for LightCycler technical assistance and Edward Tsyrlin for helping with taking mosquito photos. We also thank Chris Paton, Michael Townsend and Kyran Staunton in James Cook University for collecting field population, providing space and facility support for mosquito rearing and semi-field cage experiments.

