## Supplementary information for "Measuring the host-seeking ability of *Aedes aegypti* destined for field release"

**Supplementary Table 1.** Data of the release times, temperatures at the release point and in the Queenslander (Qld) section of the semi-field cage for each experimental replicate.

| **Experiment** | **Replicate** | **Time and date** | **Mean temperature in Qld (°C)** | **Mean temperature at release point (°C)** |
| --- | --- | --- | --- | --- |
| *Wolbachia* comparison (5 d old) | a (wing length) | 16:00 16/11/18 | 30.1 | 30.2 |
|  | b | 10:30 17/11/18 | 30.2 | 28.0 |
|  | c | 15:00 18/11/18 | 30.2 | 30.8 |
|  | d | 11:00 19/11/18 | 28.5 | 30.6 |
|  | e | 15:00 19/11/18 | 30.1 | 30.5 |
|  | f | 09:15 20/11/18 | 26.6 | 26.2 |
|  | g (wing length) | 10:00 16/11/18 | 28.0 | 27.2 |
| *Wolbachia* comparison (20 d old) | a | 09:30 24/11/18 | 29.4 | 30.4 |
|  | b | 14:00 25/11/18 | 36.5 | 34.5 |
|  | c | 09:00 29/11/18 | 31.8 | 33.6 |
|  | d | 16:00 30/11/18 | 32.2 | 32.3 |
| Laboratory maintenance comparison | a | 11:30 11/11/18 | 26.9 | 27.8 |
|  | b | 09:30 14/11/18 | 26.2 | 24.0 |
|  | c | 12:00 15/11/18 | 27.6 | 29.4 |
|  | d | 11:15 20/11/18 | 28.9 | 30.8 |
|  | e | 16:10 20/11/18 | 29.2 | 28.8 |
|  | f | 12:30 21/11/18 | 28.9 | 28.9 |
| Male test | a | 16:00 22/11/18 | 30.0 | 30.2 |

**Supplementary Table 2.** Data for average and median time to landing and total proportion landing after 42 minutes for each experimental replicate of the *Wolbachia* infection comparison.

| **Population** | **Replicate** | **Average time to landing** | **Median time to landing** | **Proportion landed** |
| --- | --- | --- | --- | --- |
| **Uninfected** | a | 6.48 | 4.5 | 0.82 |
|  | b | 8.83 | 4.5 | 0.54 |
|  | c | 7.85 | 1.5 | 0.68 |
|  | d | 10.25 | 4.5 | 0.48 |
|  | e | 10.97 | 4.5 | 0.76 |
|  | f | 12.63 | 7.5 | 0.62 |
| ***w*AlbB** | a | 7.50 | 1.5 | 0.80 |
|  | b | 7.60 | 1.5 | 0.62 |
|  | c | 6.81 | 4.5 | 0.70 |
|  | d | 7.14 | 1.5 | 0.66 |
|  | e | 6.88 | 4.5 | 0.78 |
|  | f | 9.05 | 4.5 | 0.62 |
| ***w*Mel** | a | 9.70 | 1.5 | 0.82 |
|  | b | 7.00 | 1.5 | 0.48 |
|  | c | 6.16 | 1.5 | 0.58 |
|  | d | 8.78 | 4.5 | 0.66 |
|  | e | 6.17 | 1.5 | 0.72 |
|  | f | 7.80 | 4.5 | 0.80 |

**Supplementary Table 3.** Data for average and median time to landing and total proportion landing after 42 minutes for each experimental replicate of the laboratory maintenance comparison.

| **Population** | **Replicate** | **Average time to landing** | **Median time to landing** | **Proportion landed** |
| --- | --- | --- | --- | --- |
| **Field** | a | 15.07 | 10.5 | 0.42 |
|  | b | 13.19 | 10.5 | 0.58 |
|  | c | 14.32 | 10.5 | 0.66 |
|  | d | 9.67 | 1.5 | 0.58 |
|  | e | 13.03 | 7.5 | 0.64 |
|  | f | 8.50 | 4.5 | 0.66 |
| **Laboratory** | a | 15.00 | 13.5 | 0.56 |
|  | b | 7.10 | 4.5 | 0.60 |
|  | c | 11.61 | 4.5 | 0.70 |
|  | d | 12.17 | 4.5 | 0.54 |
|  | e | 9.90 | 7.5 | 0.60 |
|  | f | 7.06 | 1.5 | 0.68 |

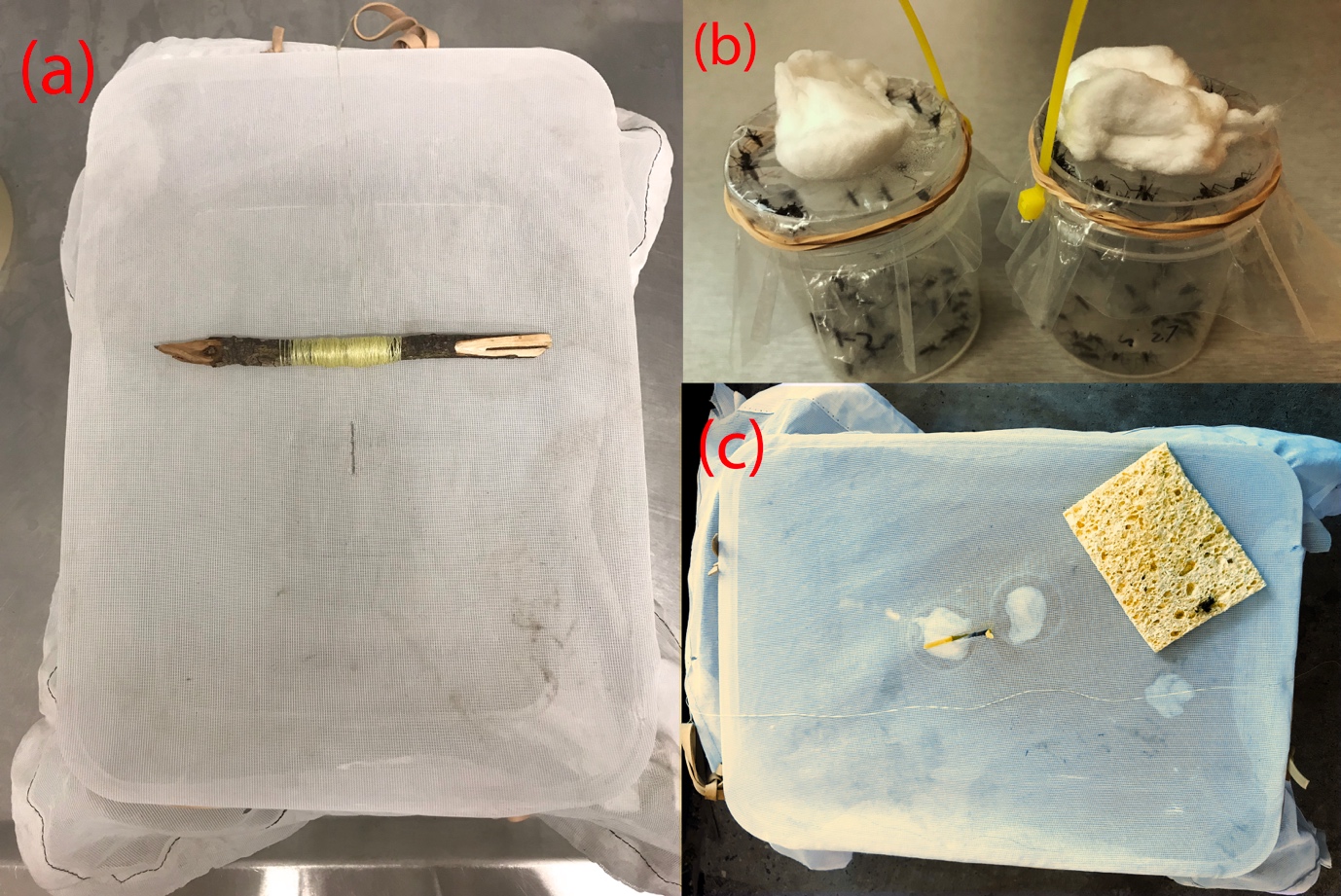

**Supplementary Figure 1.** Release mechanism. (a) Before the release, mosquitoes were trapped in the “release box” with a mesh lid that can be pulled down by a fishline from a far distance. The hole on the mesh allowed mosquitoes to be transferred and was sealed by tape; (b) Mosquitoes were marked by adding them to 70 mL specimen cups before the release, with cotton balls used to cover holes in the plastic cover that allow for mosquito transfer; (c) cable tie that can release the marked mosquitoes in the “release box” before the release to ensure different colonies of mosquitoes will be released at the same time.

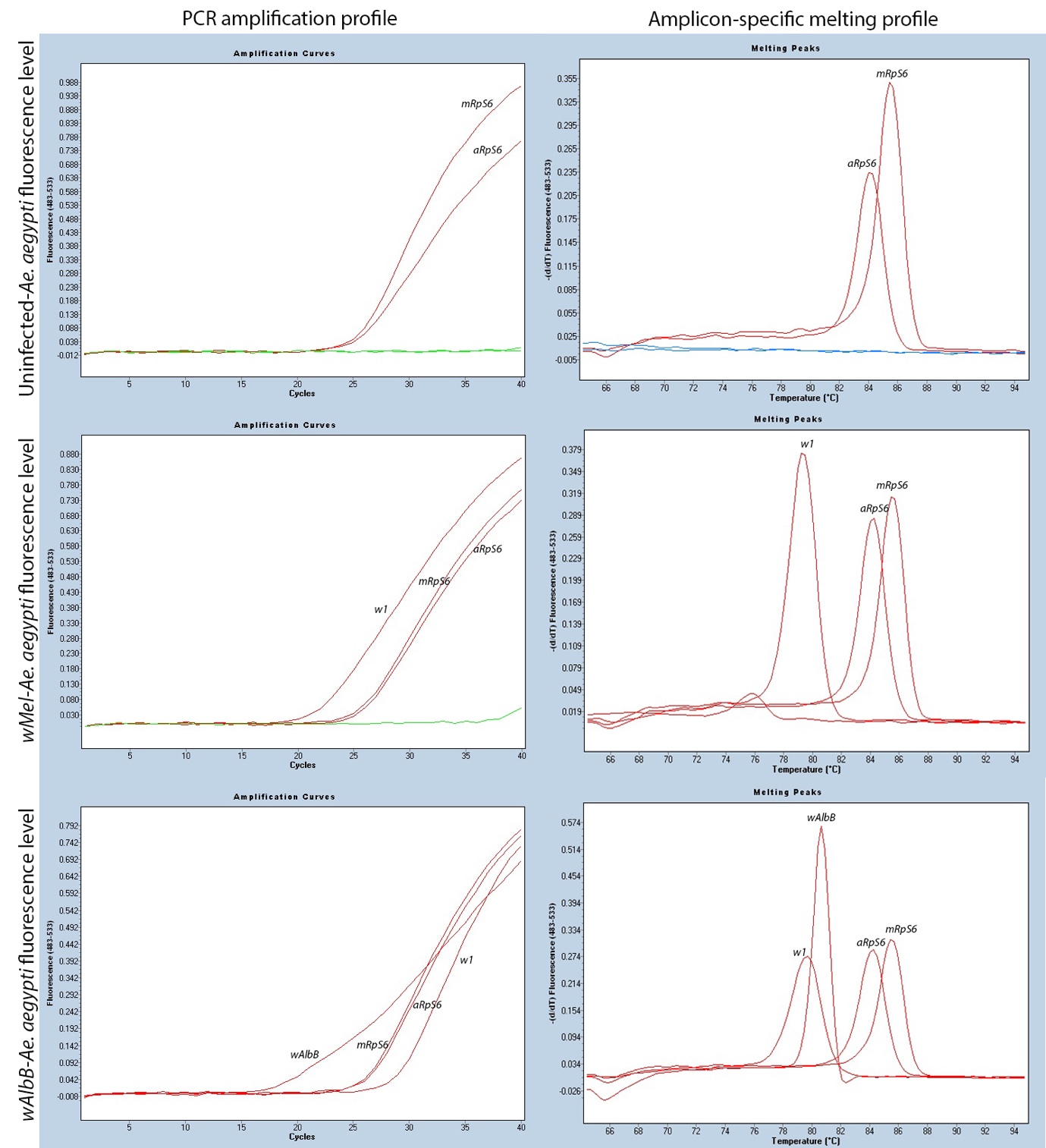

**Supplementary Figure 2.** PCR amplification profile and amplicon-specific melting profile of uninfected*, w*Mel-infected and *w*AlbB-infected *A. aegypti* using mosquito-specific (*mRpS6*) primers, *A. aegypti*-specific (*aRpS6*) primers, *Wolbachia w*Mel-specific (*w1*) primers and *Wolbachia w*AlbB-specific (*w*AlbB) primers.

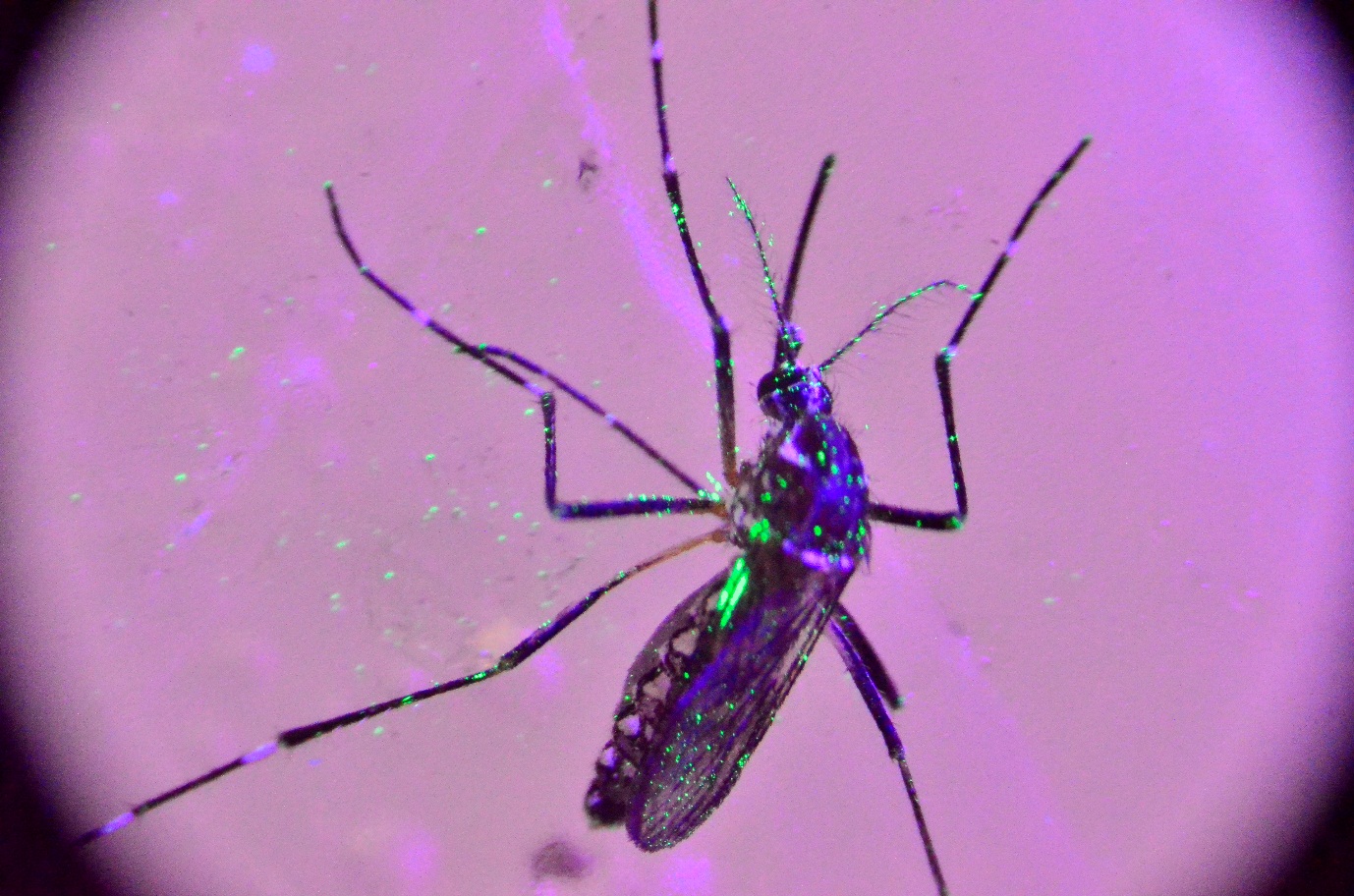

**Supplementary Figure 3.** Representative female *A. aegypti* marked with yellow fluorescent powder and visualized under UV light, demonstrating the amount of powder used for marking in the laboratory maintenance experiment.

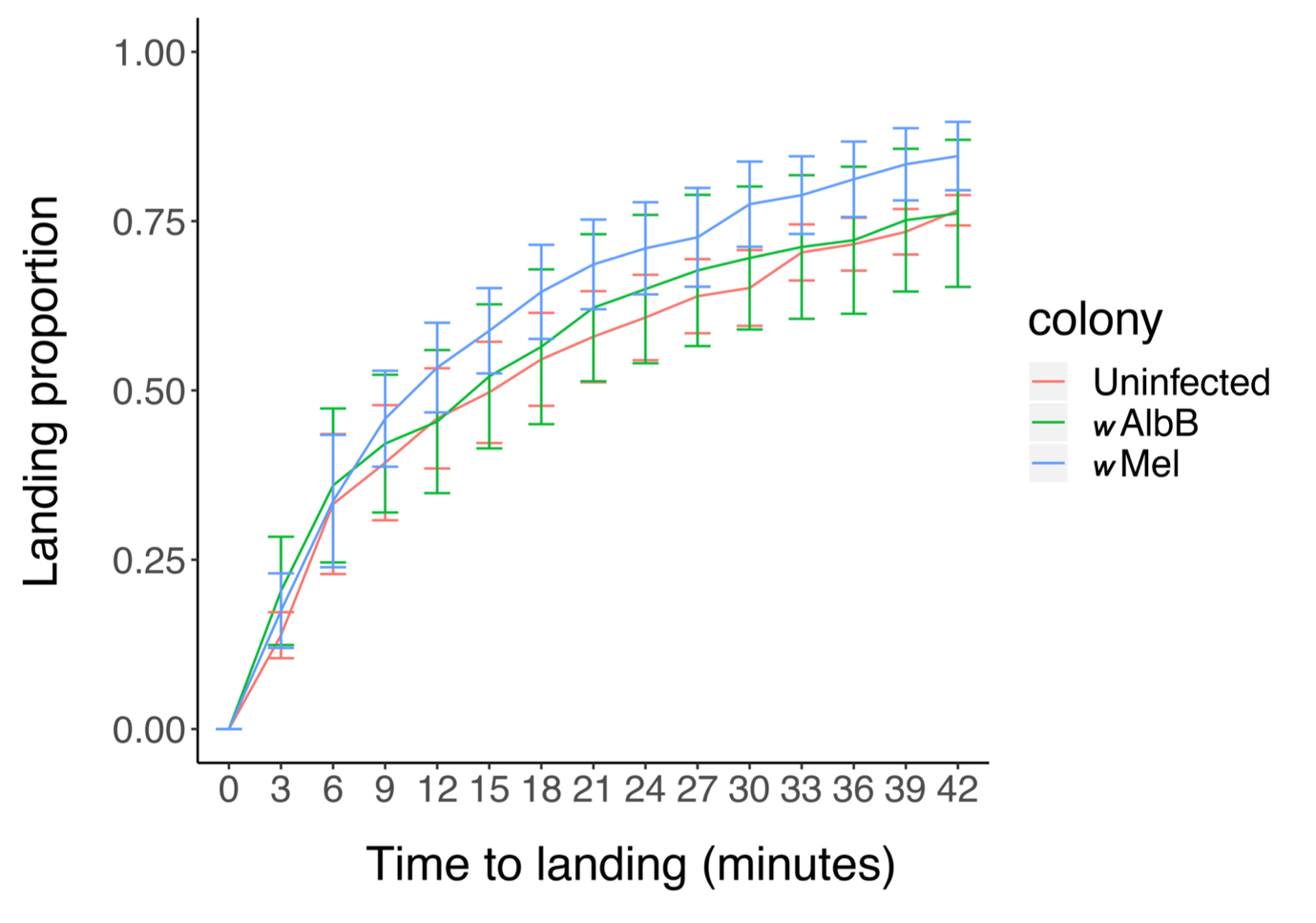

**Supplementary Figure 4.** Host-seeking ability of 20 d old *w*Mel-infected, *w*AlbB-infected and uninfected *A. aegypti* females in a semi-field cage. Cumulative landing proportions of females on human experimenters across four replicates are shown. Average arrival times were 11.4 minutes for uninfected, 10.0 minutes for *w*AlbB-infected and 10.7 minutes for *w*Mel-infected females. The proportion of females landing across experiments were 0.765 for uninfected, 0.750 for *w*AlbB-infected and 0.845 for *w*Mel-infected females*.* Lines represent means and error bars represent standard errors.

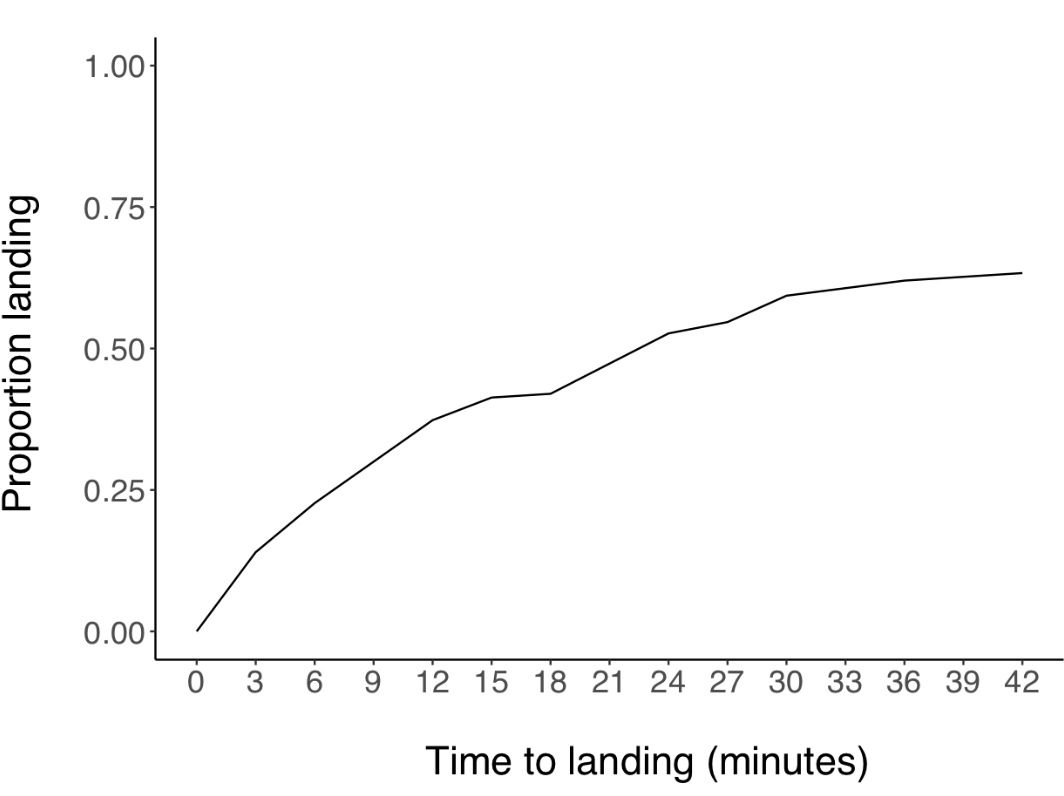

**Supplementary Figure 5.** Results of a pilot experiment testing the host-seeking ability of male *A. aegypti*. We released 150 males at the door end of the semi-field cage with an identical method to the female experiments. Two experimenters stood on a tarp under the Qld with legs exposed. Males that swarmed around the experimenters were knocked down with an electrified mosquito racket and collected with a mechanical aspirator at 3-minute intervals. Male *A. aegypti* exhibited similar host-seeking ability to females and had an average arrival time of 12.7 minutes, with 65% of the released males collected within 42 minutes.
